# Statistical test with sample size and power calculation for paired repeated measures designs of method comparison studies

**DOI:** 10.1101/516658

**Authors:** Yun Bai, Zengri Wang, Theodore C. Lystig, Baolin Wu

**Affiliations:** Medtronic plc, Minneapolis, MN, USA; Division of Biostatistics, School of Public Health, University of Minnesota

**Keywords:** Paired repeated measures, Pulse oximetry studies, Sample size and power calculation

## Abstract

The paired repeated measures (PRM) design has been commonly used in comparison studies to demonstrate that two measures agree sufficiently up to a pre-specified threshold. Due to the nature of dependence in repeated measures, existing methods for analyzing these comparison studies are often ad hoc and less than satisfactory. We propose a new test together with sample size and power calculation for PRM designs, which exhibit very good finite sample performance. We demonstrate favorable performance of the proposed methods through numerical studies. We provide very efficient R programs for our methods in a publicly available R package.

## 1 Introduction

Paired repeated measures designs of method comparison studies are commonly used to demonstrate the effectiveness of new measures relative to a standard measure in areas such as pulse oximetry studies. An oximeter is a device used to transmit radiation at a known wavelength(s) through blood to measure the blood oxygen saturation based on the amount of reflected or scattered radiation. It may be used alone or in conjunction with a fiberoptic oximeter catheter. These studies often set out to test whether two measures agree sufficiently with respect to some pre-specified agreement score, e.g., the commonly used average root mean squared error (*A*_*RMS*_). Some early developed methods for analyzing such comparison studies are mostly descriptive. For example, a commonly used measure is the concordance correlation coefficient, which measures the agreement between two variables to evaluate reproducibility or for inter-rater reliability (Lin, 1989, 1992). The Bland-Altman plot is another popular exploratory analysis approach (Bland and Altman, 1999, 2007). In engineering applications, a measurement error model known as Deming regression (Deming, 1964), which allows for errors in both the predictor and response variables, is commonly used.

Recently some formal hypothesis testing methods have been proposed (Pennello, 2002, 2003; Ndikintum and Rao, 2016). They are based on the asymptotic normal distribution approximation and do not have good finite sample performance in our numerical studies. Furthermore the corresponding sample size and power calculation methods accompanying these hypothesis testing solutions are not well studied and generally ad hoc with unsatisfactory performance.

The purpose of this research practice note is to rigorously characterize the finite sample distribution of the *A*_*RMS*_ and develop efficient and powerful test methods together with corresponding sample size and power calculation methods. We will demonstrate the utility of our proposed methods through thorough numerical studies, and show that they perform very favorably compared to the existing methods. We further develop efficient numerical algorithms and implement them in a publicly available R package. Our proposed methods and developed R package provide practical tools which bridge an existing gap in the field. Throughout the paper, we will mainly focus on intuitive ideas and delegate all technical details to the Appendix.

## 2 Statistical methods

For unit *i* = 1,2, ⋯, *n* and paired measurements *j* = 1,2, ⋯,*m*_*i*_ for unit *i*, let *d*_*ij*_ = *y*_*ij*_ − *x*_*ij*_ denote the difference between the paired measures *y*_*ij*_ and *x*_*ij*_ for the *ij*-th pair (e.g., pulse oximeter measurement and the co-oximeter measurement). Following previous approaches, we model *d*_*ij*_ = *μ* + *μ* + *ϵ*_*ij*_, where 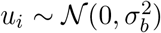 and 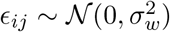. Here 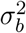 is the between-unit variance, 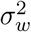 is the within-unit variance, and *μ* quantifies the average measure difference. Let *D_i_* × (*d_i1_*; ⋯, *d_im_i__*)^*T*^. We can readily compute its covariance matrix σ_*i*_ = *Cov*(*D*_*i*_) 
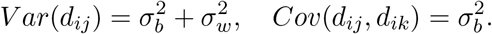

Let 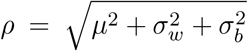. The paired repeated measure (PRM) comparison is based on testing *H*_0_: *ρ* ≥ *ρ*_0_ versus *H*_*a*_: *ρ* < *ρ*_0_, where *ρ*_0_ is a pre-specified acceptable threshold. For example, the FDA regards a pulse oximeter as approvable if the null hypothesis *H*_0_: *ρ* ≥ *ρ*_0_ can be rejected in favor of the alternative hypothesis *H*_a_: *ρ* < *ρ*_0_ with, e.g., *ρ*_0_ = 3% (see https://www.fda.gov/RegulatoryInformation/Guidances/ucm341718.htm).

### 2.1 Existing methods

Let 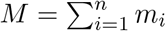. A direct measure of the accuracy is the root mean square defined as 
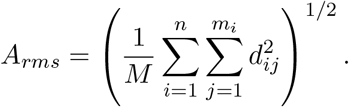

It is challenging to derive the null distribution of *A*_*rms*_. Traditionally the bootstrap approach has been used to compute the variation of *A*_*rms*_, which is then used to conduct hypothesis testing based on the large sample normal approximation. Sample size and power calculation is even more challenging due to the need to derive both the null and alternative distributions.

An efficient approach is due to Pennello (2002, 2003). He recognized the quadratic form of 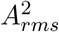 and derived its analytical mean and variance, which are then used to form a large sample normal approximation to compute the sample size and power for a PRM study. Specifically note that 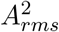 is constructed using a sum of *i.i.d.* terms. Hence we can approximately model it using a normal distribution 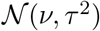, where *v* and *τ*^2^ are estimated by matching the first two moments. For example, when *m*_*i*_ = *m*, we can easily check that 
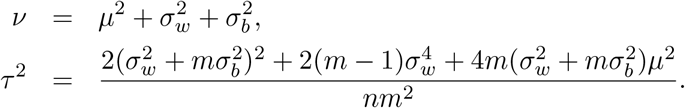

An approximate Z-test based on a large sample normal approximation can then be constructed 
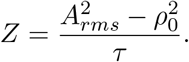

Under the null, asymptotically *Z* is distributed as 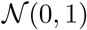 and generally 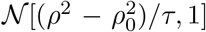. This can be used to perform hypothesis testing and sample size/power calculations. Ndikintum and Rao (2016) further generalize Pennello’s normal approximation method based on the score and Wald based Z-tests (denoted as Z-score and Z-Wald in the following discussion).

We note that the finite sample performance might be very poor for these normal approximation based methods. It might need a very large sample size *n* for the asymptotic normal distribution to kick in. Next we discuss our proposed methods, which are based on maximum likelihood estimation, and rely on the analytical finite-sample distribution without the need for large sample approximation. Our proposed methods can readily handle any balanced or unbalanced designs, and demonstrate very good performance in numerical studies.

### 2.2 Parameter estimation: MLE and REML

We first discuss the efficient calculation of log likelihood for the PRM data. Given 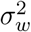 and 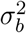 we can analytically compute the inverse and determinant of Σ_*i*_ as 
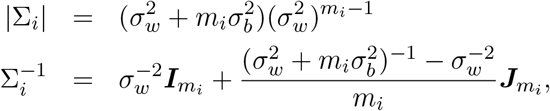
where ***I***_*N*_ denotes the *N*-th order identity matrix and ***J***_*N*_ an *N* × *N* matrix of ones. Hence we can analytically compute −2 log[Pr(*D*_*i*_)] as (up to some constant that does not depend on data) 
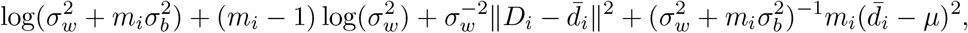
where 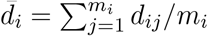. The maximum likelihood estimate (MLE) can be obtained by maximizing the log likelihood 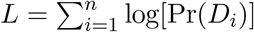. To reduce the bias of variance parameter estimates, we can use the restricted maximum likelihood (REML) approach, which can be checked to be based on maximizing 
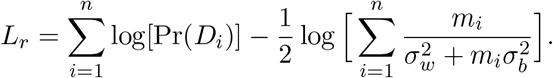

### 2.3 PRM comparison: hypothesis testing

We propose to base the PRM comparison on the test statistic 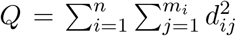. One can readily check that 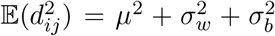, and hence 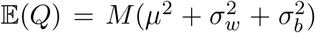, where 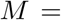 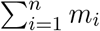.

Given 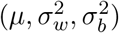, we can analytically show that *Q* is distributed as the weighted sum of independent chi-square random variables, 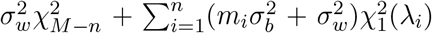, where 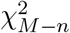 denotes a central chi-square random variable with (*M* − *n*) degrees of freedom (DF), 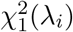 denotes an 1-DF non-central chi-square random variable with non-centrality parameter 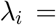 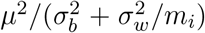. For balanced designs with 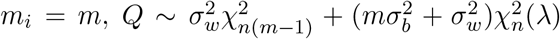, where the non-centrality parameter 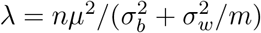.

With the derived analytical distribution for *Q*, we can readily compute its p-value as follows. We compute the test statistic *Q* and obtain the parameter estimates, 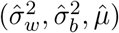, under the null based on the MLE or REML. We then refer *Q* to the distribution, 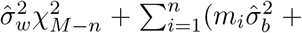 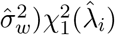, to compute the significance p-value using the Davies method (Davies, 1980).

### 2.4 Sample size and power calculation

Given the true parameter values 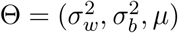 and the accepted null hypothesis threshold 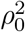, we want to evaluate the power for a given sample size (*n*, *m*). The analytical power calculation for the PRM design problem is not trivial due to its nature of composite hypothesis testing: we are testing a quadratic sum of parameter values. Existing methods have made simplified assumptions and computed power only under pre-specified null parameter values.

One straightforward though computing-intensive approach is based on Monte Carlo simulation. At the *b*-th simulation, we generate the data based on the PRM design under the given parameter values as follows. First we simulate independent standard normal random numbers, (*x*_*ij*_,*u*_*i*_), *i* = 1, ⋯,*n*, *j* = 1, ⋯,*m*. We then define the data *d*_*ij*_ = *x*_*ij*_σ_*w*_ + *u*_*i*_σ_*b*_ + *μ*. Given the simulated data, we can compute the test p-value, denoted as *p*_*b*_, based on the proposed method. Over *B* Monte Carlo simulations, we can evaluate our power at significance level *α* as 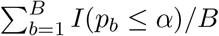, where *I*() is the indicator function.

Alternatively we propose an analytical approach based on the average log likelihood as follows. Note that the key to power calculation is to solve the quantile of a test statistic *Q* at the given a level under the null hypothesis. This requires the null parameter values 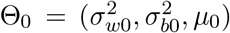, which are estimated as follows. Instead of generating data from Monte Carlo simulation and defining the log likelihood to estimate parameters repeatedly, we average the log likelihood over Monte Carlo simulations to estimate parameters. And it turns out we can analytically compute the average log likelihood without actual Monte Carlo simulation. Specifically we define *L*(Θ_0_) = 𝔼{log[Pr(*D*|Θ_0_)]|Θ}, where the data *D* = (*d*_1_ ⋯, *d*_*m*_)^*T*^ are generated under the PRM design assuming parameters Θ, the inner log likelihood log[Pr(*D*|Θ_0_)] is computed assuming parameters Θ_0_, and the outside expectation is taken assuming parameters Θ. One can check that *L*(Θ_0_) is essentially the average log likelihood over Monte Carlo simulations. We then estimate 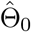 based on maximizing the average log likelihood, 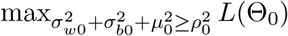. We compute the quantile of the test statistic at significance level *α*, denoted as *q*_*α*_, assuming parameters 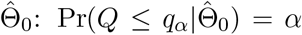. Lastly we compute power as Pr(*Q* ≤ *q*_α_|Θ). Both probabilities are calculated using the Davies method as shown previously.

It can be shown that −2L(Θ_0_) is equal to (up to some constant; see Appendix) 
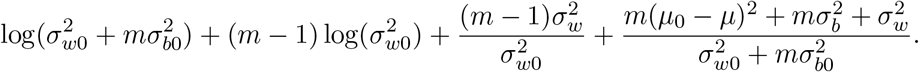

Statistically this average log likelihood approach amounts to minimizing the Kullback-Leibler divergence of the hypothesized null distribution from the true data generating distribution.

Similarly we can take a REML approach by maximizing 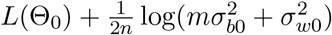.

## 3 Numerical studies

In the numerical studies below, we use REML for illustration. Both MLE and REML are implemented in our publicly available R package. MLE leads to very similar results.

### 3.1 Type I errors

To evaluate type I errors, we consider 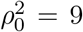 and the combinations of 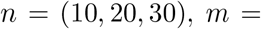 with 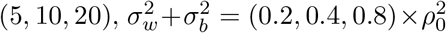, and setting 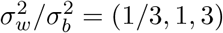.

We conduct 10^5^ null simulations to evaluate the type I errors at the significance level *α* = 0.001, 0.01 and 0.05. Table 1 shows the empirical type I errors for 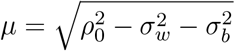 and 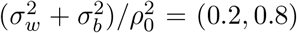 and *α* = 0.01. Similar results have been observed for other settings. The proposed test (denoted as QMS) is compared to the Z-score and Z-Wald tests. Overall the results show that the type I errors are well controlled for the proposed method, while the Z-score test is very conservative and Z-Wald test has largely inflated type I errors. For the proposed QMS, (1) not surprisingly, larger *n* leads to more accurate control of type I errors; (2) in contrast, QMS has very similar type I error control across *m*, the number of repeated measures; (3) generally smaller within-subject variance 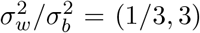 also leads to slightly more accurate control of type I errors; and (4) larger *μ* (smaller 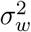) also leads to more accurate control of type I errors.

**Table 1:**
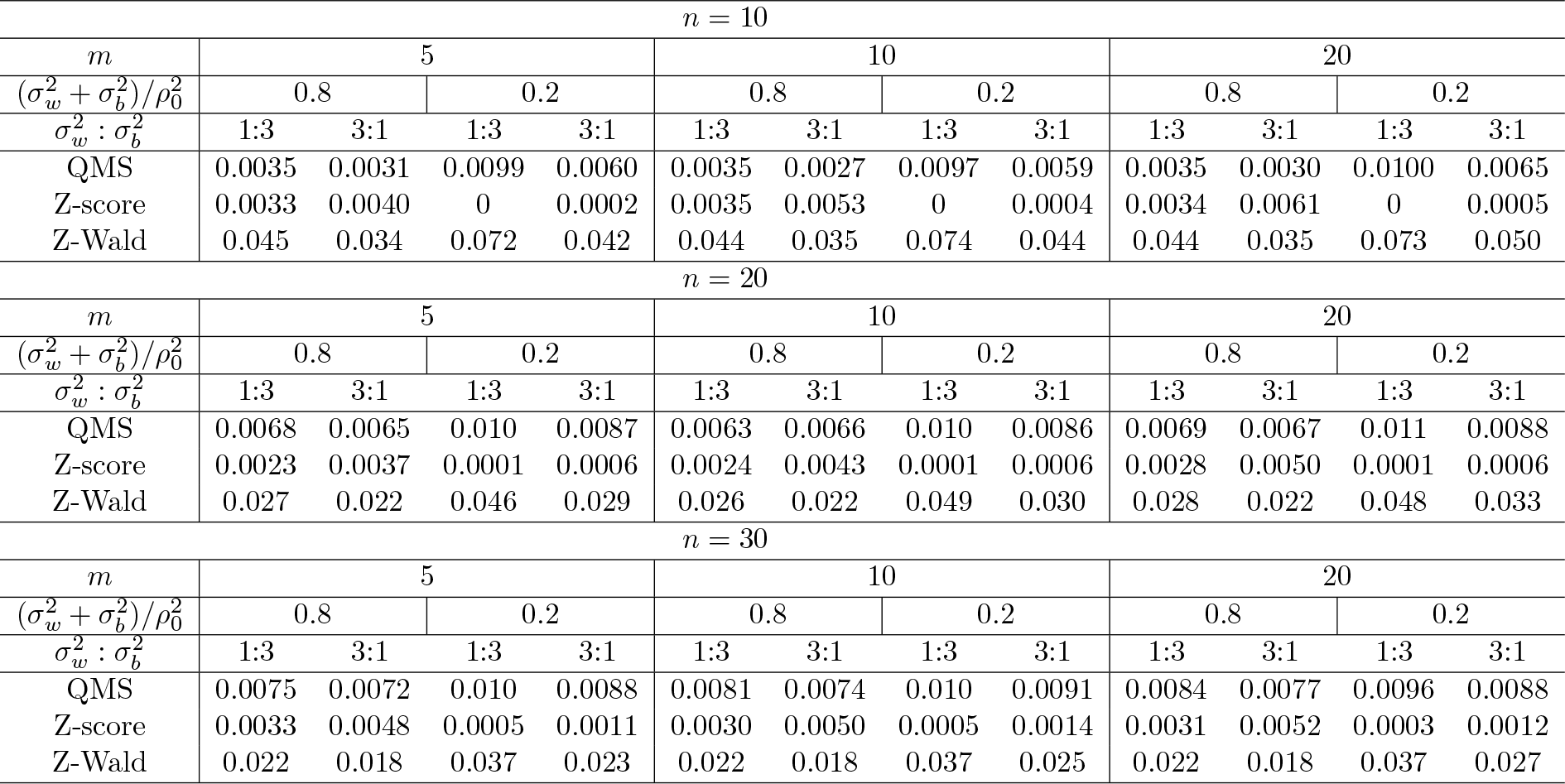
Empirical type I errors at 0.01 significance level estimated over 10^5^ null simulations. Data are simulated from *n* subjects each with *m* measures. We are testing 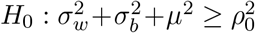. QMS is the proposed test, Z-score and Z-Wald are the score and Wald based Z-tests.

| $n = 10$ | | | | | | | | | | | | |
| --- | --- | --- | --- | --- | --- | --- | --- | --- | --- | --- | --- | --- |
| $m$ | 5 | | | | 10 | | | | 20 | | | |
| $(\sigma_w^2 + \sigma_b^2)/\rho_0^2$ | 0.8 | | 0.2 | | 0.8 | | 0.2 | | 0.8 | | 0.2 | |
| $\sigma_w^2 : \sigma_b^2$ | 1:3 | 3:1 | 1:3 | 3:1 | 1:3 | 3:1 | 1:3 | 3:1 | 1:3 | 3:1 | 1:3 | 3:1 |
| QMS | 0.0035 | 0.0031 | 0.0099 | 0.0060 | 0.0035 | 0.0027 | 0.0097 | 0.0059 | 0.0035 | 0.0030 | 0.0100 | 0.0065 |
| Z-score | 0.0033 | 0.0040 | 0 | 0.0002 | 0.0035 | 0.0053 | 0 | 0.0004 | 0.0034 | 0.0061 | 0 | 0.0005 |
| Z-Wald | 0.045 | 0.034 | 0.072 | 0.042 | 0.044 | 0.035 | 0.074 | 0.044 | 0.044 | 0.035 | 0.073 | 0.050 |
| $n = 20$ | | | | | | | | | | | | |
| $m$ | 5 | | | | 10 | | | | 20 | | | |
| $(\sigma_w^2 + \sigma_b^2)/\rho_0^2$ | 0.8 | | 0.2 | | 0.8 | | 0.2 | | 0.8 | | 0.2 | |
| $\sigma_w^2 : \sigma_b^2$ | 1:3 | 3:1 | 1:3 | 3:1 | 1:3 | 3:1 | 1:3 | 3:1 | 1:3 | 3:1 | 1:3 | 3:1 |
| QMS | 0.0068 | 0.0065 | 0.010 | 0.0087 | 0.0063 | 0.0066 | 0.010 | 0.0086 | 0.0069 | 0.0067 | 0.011 | 0.0088 |
| Z-score | 0.0023 | 0.0037 | 0.0001 | 0.0006 | 0.0024 | 0.0043 | 0.0001 | 0.0006 | 0.0028 | 0.0050 | 0.0001 | 0.0006 |
| Z-Wald | 0.027 | 0.022 | 0.046 | 0.029 | 0.026 | 0.022 | 0.049 | 0.030 | 0.028 | 0.022 | 0.048 | 0.033 |
| $n = 30$ | | | | | | | | | | | | |
| $m$ | 5 | | | | 10 | | | | 20 | | | |
| $(\sigma_w^2 + \sigma_b^2)/\rho_0^2$ | 0.8 | | 0.2 | | 0.8 | | 0.2 | | 0.8 | | 0.2 | |
| $\sigma_w^2 : \sigma_b^2$ | 1:3 | 3:1 | 1:3 | 3:1 | 1:3 | 3:1 | 1:3 | 3:1 | 1:3 | 3:1 | 1:3 | 3:1 |
| QMS | 0.0075 | 0.0072 | 0.010 | 0.0088 | 0.0081 | 0.0074 | 0.010 | 0.0091 | 0.0084 | 0.0077 | 0.0096 | 0.0088 |
| Z-score | 0.0033 | 0.0048 | 0.0005 | 0.0011 | 0.0030 | 0.0050 | 0.0005 | 0.0014 | 0.0031 | 0.0052 | 0.0003 | 0.0012 |
| Z-Wald | 0.022 | 0.018 | 0.037 | 0.023 | 0.022 | 0.018 | 0.037 | 0.025 | 0.022 | 0.018 | 0.037 | 0.027 |

### 3.2 Power

Since the Z-Wald test has severely inflated type I errors, we only include the proposed test and the Z-score test in the power comparison. We consider the combinations of *n* = (10,20,30), 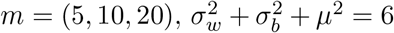 with 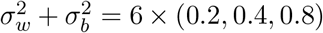 and 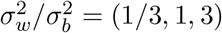. We test 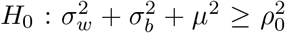 with *ρ*_0_ = 3. Generally larger 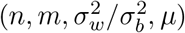 will lead to larger rejection power. Here we note that the estimation of between-unit variance 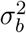 generally has larger variation compared to the within-unit variance 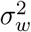, since we are essentially having sample size *n* and *n*(*m* − 1) respectively for these two variance parameters (see Appendix). Hence larger 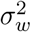 and smaller 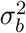 will lead to relatively more accurate estimation and hence larger rejection power.

We use 10^4^ Monte Carlo simulations to estimate power under each configuration. Table 2 summarizes the power for 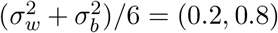 and 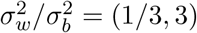 under 0.05 significance level. Similar patterns are observed for other settings. Overall the proposed method performs better than the Z-score test by a large margin. For the proposed QMS, (1) not surprisingly, increasing either *n* or *m* can lead to larger power, and generally increasing n brings more power improvement; (2) smaller between-subject variance 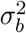 leads to larger power; and (3) larger *μ* (smaller 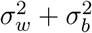) also leads to larger power.

**Table 2:** Power (%) at 0.05 significance level estimated over 10^4^ simulations. Data are simulated for n subjects each with *m* measures under 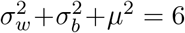. We are testing 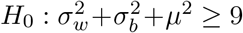. QMS is the proposed test, and Z-score is the score based Z-tests.

| $n = 10$ | | | | | | | | | | | | |
| --- | --- | --- | --- | --- | --- | --- | --- | --- | --- | --- | --- | --- |
| $m$ | 5 | | | | 10 | | | | 20 | | | |
| $(\sigma_w^2 + \sigma_b^2)/6$ | 0.8 | | 0.2 | | 0.8 | | 0.2 | | 0.8 | | 0.2 | |
| $\sigma_w^2 : \sigma_b^2$ | 1:3 | 3:1 | 1:3 | 3:1 | 1:3 | 3:1 | 1:3 | 3:1 | 1:3 | 3:1 | 1:3 | 3:1 |
| QMS | 23.5 | 39.6 | 42.4 | 68.6 | 23.5 | 47.7 | 43.1 | 78.2 | 24.1 | 54.3 | 44.3 | 82.2 |
| Z-score | 1.0 | 5.2 | 11.2 | 31.6 | 1.2 | 6.8 | 11.6 | 41.6 | 1.0 | 8.1 | 12.1 | 47.0 |
| $n = 20$ | | | | | | | | | | | | |
| $m$ | 5 | | | | 10 | | | | 20 | | | |
| $(\sigma_w^2 + \sigma_b^2)/\rho_0^2$ | 0.8 | | 0.2 | | 0.8 | | 0.2 | | 0.8 | | 0.2 | |
| $\sigma_w^2 : \sigma_b^2$ | 1:3 | 3:1 | 1:3 | 3:1 | 1:3 | 3:1 | 1:3 | 3:1 | 1:3 | 3:1 | 1:3 | 3:1 |
| QMS | 39.0 | 68.7 | 75.9 | 96.0 | 41.5 | 80.8 | 76.3 | 98.5 | 42.4 | 86.2 | 77.3 | 99.3 |
| Z-score | 12.0 | 37.3 | 51.0 | 86.7 | 13.1 | 56.1 | 53.7 | 96.3 | 12.8 | 49.3 | 53.0 | 93.8 |
| $n = 30$ | | | | | | | | | | | | |
| $m$ | 5 | | | | 10 | | | | 20 | | | |
| $(\sigma_w^2 + \sigma_b^2)/\rho_0^2$ | 0.8 | | 0.2 | | 0.8 | | 0.2 | | 0.8 | | 0.2 | |
| $\sigma_w^2 : \sigma_b^2$ | 1:3 | 3:1 | 1:3 | 3:1 | 1:3 | 3:1 | 1:3 | 3:1 | 1:3 | 3:1 | 1:3 | 3:1 |
| QMS | 54.6 | 85.7 | 90.6 | 99.4 | 56.3 | 93.2 | 91.2 | 99.92 | 56.7 | 96.7 | 92.0 | 99.97 |
| Z-score | 25.3 | 64.3 | 77.0 | 98.1 | 27.2 | 77.3 | 79.2 | 99.45 | 27.9 | 85.6 | 79.7 | 99.77 |

In addition, we also evaluate the power calculation. We consider the same simulation settings as previously and treat the Monte Carlo results as our gold standard. We then apply the proposed analytical power calculation method and compare those results to the gold standard results from the Monte Carlo simulations. Table 3 summarizes the results for 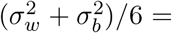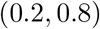 and 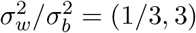. Overall we have observed that the analytically computed powers (denoted as ACP) provide a good approximation to the Monte Carlo results (denoted as MCP). The proposed analytical power calculation (ACP) generally performs well under larger sample size *n* and smaller *μ*.

**Table 3:** Computed power (%) at 0.05 significance level for the proposed test QMS: Monte Carlo results are displayed as MCP, while the analytically computed powers are displayed as ACP. Data are simulated for *n* subjects each with *m* measures under 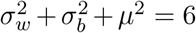. We are testing 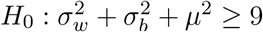.

| $n = 10$ | | | | | | | | | | | | |
| --- | --- | --- | --- | --- | --- | --- | --- | --- | --- | --- | --- | --- |
| $m$ | 5 | | | | 10 | | | | 20 | | | |
| $(\sigma_w^2 + \sigma_b^2)/6$ | 0.8 | | 0.2 | | 0.8 | | 0.2 | | 0.8 | | 0.2 | |
| $\sigma_w^2 : \sigma_b^2$ | 1:3 | 3:1 | 1:3 | 3:1 | 1:3 | 3:1 | 1:3 | 3:1 | 1:3 | 3:1 | 1:3 | 3:1 |
| MCP | 23.5 | 39.6 | 42.4 | 68.6 | 23.5 | 47.7 | 43.1 | 78.2 | 24.1 | 54.3 | 44.3 | 82.2 |
| ACP | 21.4 | 36.7 | 39.2 | 61.1 | 22.0 | 43.8 | 40.1 | 68.4 | 22.3 | 48.9 | 40.5 | 72.7 |

| $n = 20$ | | | | | | | | | | | | |
| --- | --- | --- | --- | --- | --- | --- | --- | --- | --- | --- | --- | --- |
| $m$ | 5 | | | | 10 | | | | 20 | | | |
| $(\sigma_w^2 + \sigma_b^2)/\rho_0^2$ | 0.8 | | 0.2 | | 0.8 | | 0.2 | | 0.8 | | 0.2 | |
| $\sigma_w^2 : \sigma_b^2$ | 1:3 | 3:1 | 1:3 | 3:1 | 1:3 | 3:1 | 1:3 | 3:1 | 1:3 | 3:1 | 1:3 | 3:1 |
| QMS | 39.0 | 68.7 | 75.9 | 96.0 | 41.5 | 80.8 | 76.3 | 98.5 | 42.4 | 86.2 | 77.3 | 99.3 |
| Z-score | 38.5 | 66.1 | 71.2 | 92.7 | 39.7 | 76.5 | 72.5 | 96.3 | 40.3 | 82.5 | 73.1 | 97.7 |

| $n = 30$ | | | | | | | | | | | | |
| --- | --- | --- | --- | --- | --- | --- | --- | --- | --- | --- | --- | --- |
| $m$ | 5 | | | | 10 | | | | 20 | | | |
| $(\sigma_w^2 + \sigma_b^2)/\rho_0^2$ | 0.8 | | 0.2 | | 0.8 | | 0.2 | | 0.8 | | 0.2 | |
| $\sigma_w^2 : \sigma_b^2$ | 1:3 | 3:1 | 1:3 | 3:1 | 1:3 | 3:1 | 1:3 | 3:1 | 1:3 | 3:1 | 1:3 | 3:1 |
| MCP | 54.6 | 85.7 | 90.6 | 99.4 | 56.3 | 93.2 | 91.2 | 99.92 | 56.7 | 96.7 | 92.0 | 99.97 |
| ACP | 53.3 | 83.7 | 88.0 | 99.0 | 55.0 | 91.7 | 88.9 | 99.71 | 55.8 | 95.2 | 89.4 | 99.88 |

Given the desired power level under a pre-specified significance level, we can invert the proposed analytical power calculation method to solve the required sample size problem. Due to the discrete nature of sample size numbers, we recognize that the actual power under the computed sample size generally does not exactly equal to the desired power. But nonetheless, we expect them to provide a good approximation for design guidance. For illustration, we consider 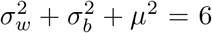 and testing 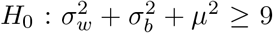. We compute sample size *n* for 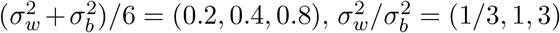 and *m* =(5,10, 20) to achieve 80% power under 0.05 significance level. Table 4 shows the calculated sample sizes together with the actual powers computed from the proposed analytical method (denoted as ACP) and 10^4^ Monte Carlo simulations (denoted as MCP). Overall we can see that the actual powers under the computed sample sizes are close to the desired levels, especially under relatively larger sample size *n*.

**Table 4:**
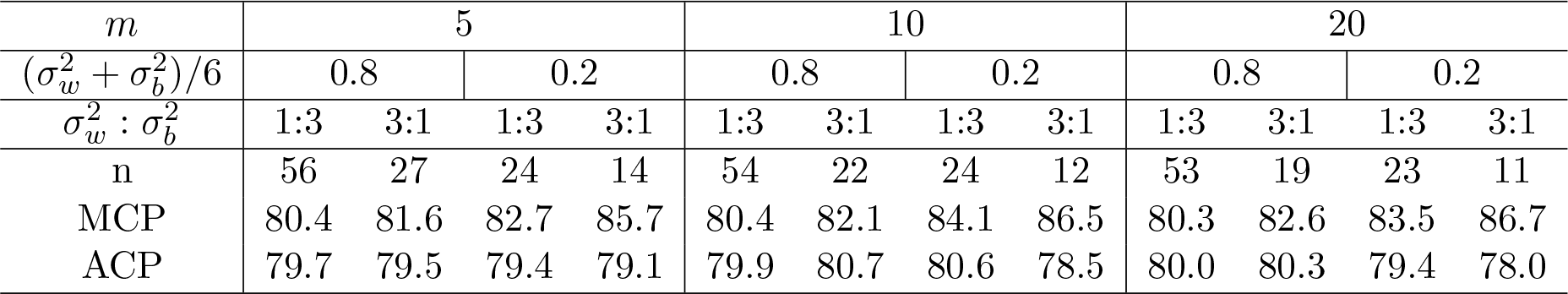
Sample size calculation to achieve 80% power at 0.05 significance level for the proposed test QMS. Also shown are the actual power computed under the calculated sample size: Monte Carlo results are displayed as MCP, while the analytically computed powers are displayed as ACP. Data are simulated for *n* subjects each with m measures under 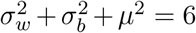. We are testing 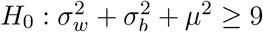.

## 4 Discussion

We have proposed a powerful test together with efficient sample size and power calculations for the paired repeated measures (PRM) designs problem. In contrast to the existing methods, which are either mostly exploratory or have relied on ad hoc large sample approximation with unsatisfactory performance, our proposed methods are based on the finite sample exact distribution and built on solid statistical principles. Numerical studies have demonstrated the very favorable performance of our proposed methods. All proposed methods have been implemented in a publicly available R package, at http://www.github.com/baolinwu/SPprm. The proposed methods and developed R package provide very useful and practical tools. Supplementary materials contain more simulation results and sample scripts to install and use the developed R package.

## 5 Acknowledgments

This research was supported in part by NIH grant GM083345 and CA134848. We are grateful to the University of Minnesota Supercomputing Institute for assistance with the computations.

## A A Likelihood calculation and parameter estimation

We can write 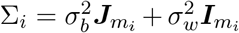. Note 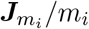 is a rank-one projection matrix. Hence Σ_*i*_ has two unique eigenvalues, 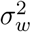 (with algebraic multiplicity *m*_*i*_ − 1) and 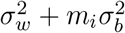 (with algebraic multiplicity 1), and we can efficiently compute its inverse and determinant as shown previously. We then analytically compute −2 log[Pr(*D*_*i*_)] as (up to some constant that does not depend on data) 
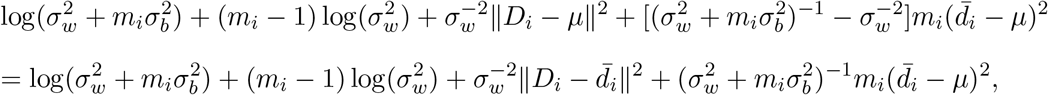
where 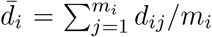. Following the approach of Laird and Ware (1982), we compute the residual log likelihood based on integrating out *μ* from the joint likelihood.

When there are no constraints on the parameters, we can easily check that 
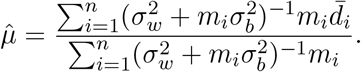

In the simple scenario of a balanced design with a constant number of observations for each unit, *m*_*i*_ = *m*, we can check that 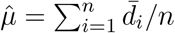.

For our proposed methods, the key is to estimate the parameters under 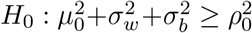, which is a constrained optimization problem solved using the Powell method (Powell, 1994).

### B Distribution of *Q*

Given 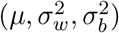, we can derive the analytical distribution of *Q* as follows. Note that we can write *d*_*ij*_ — *d̄* = *ϵ*_*ij*_ — *ϵ̄*_*i*_, where 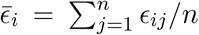, and *d̄* = *μ* + *u*_*i*_ + *ϵ̄*_*i*_. Further 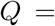 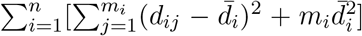. Therefore 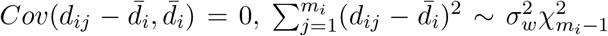 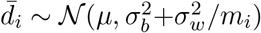. And statistically *Q* is distributed as the weighted sum of independent chi-square random variables, 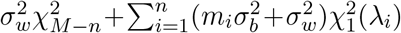, where the non-centrality parameter 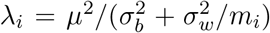. When 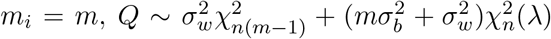, where the noncentrality parameter 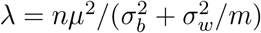.

### C Average log likelihood for power calculation

Given *D* = (*d*_1_, ⋯,*d*_*m*_)^*T*^, we can compute −2 log[Pr(*D*)|Θ_0_] as shown previously 
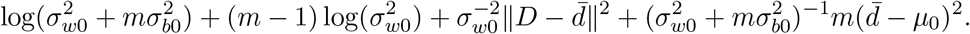

As discussed previously, 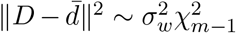, and 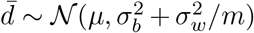. Therefore the average log likelihood −2𝔼{log[Pr(*D*|Θ_0_)]|Θ} is 
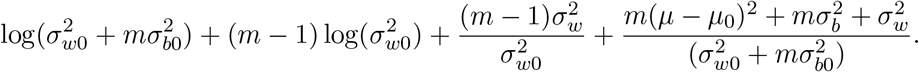

Statistically our proposed average log likelihood maximization essentially amounts to minimizing the Kullback-Leibler divergence from the hypothesized null distribution Pr(*D*|Θ_0_) to the true data generating distribution Pr(*D*|Θ) (Kullback and Leibler, 1951).

## References

Bland, J. M. and Altman, D. G. (1999). Measuring agreement in method comparison studies. Statistical Methods in Medical Research 8, 135–160.

Bland, J. M. and Altman, D. G. (2007). Agreement between methods of measurement with multiple observations per individual. Journal of Biopharmaceutical Statistics 17, 571–582.

Davies, R.B. (1980) Algorithm AS 155: the distribution of a linear combination of χ^2^ random variables. Applied Statistics, 29 (3), 323–333.

Deming, W. E. (1964). Statistical adjustment of data. Dover Publications.

Kullback, S. and Leibler, R. A. (1951). On Information and Sufficiency. The Annals of Mathematical Statistics 22, 79–86.

Laird, N. M. and Ware, J. H. (1982). Random-Effects Models for Longitudinal Data. Biometrics 38, 963–974.

Lin, L. I.-K. (1989). A Concordance Correlation Coefficient to Evaluate Reproducibility. Biometrics 45, 255–268.

Lin, L. I.-K. (1992). Assay Validation Using the Concordance Correlation Coefficient. Biometrics 48, 599–604.

Ndikintum, N. K. and Rao, M. (2016). A Special Inference Problem in Repeated Measures Design Test of Statistical Hypothesis on Accuracy Root Mean Square Application to Pulse Oximetry Studies. Statistics in Biopharmaceutical Research 8, 60–76.

Pennello, G. A. (2002). Sample size for paired repeated measures designs of method comparison studies: Application to pulse oximetry. ASA Proceedings of the Joint Statistical Meetings pages 2671–2675.

Pennello, G. A. (2003). Comparing monitoring devices when a gold standard in unavailable: Application to pulse oximeters. ASA Proceedings of the Joint Statistical Meetings pages 3256–3263.

Powell, M. J. D. (1994). A Direct Search Optimization Method That Models the Objective and Constraint Functions by Linear Interpolation. In Advances in Optimization and Numerical Analysis, Mathematics and Its Applications, pages 51–67. Springer, Dordrecht.

