## Supplementary Materials for "Statistical test with sample size and power calculation for paired repeated measures designs of method comparison studies"

#### 1 MLE simulation results

The following results are obtained based on MLE estimates.

##### 1.1 Type I errors

To evaluate type I errors, we consider the combinations of $n = (10, 20, 30)$, $m = (5, 10, 20)$, $\sigma_w^2 + \sigma_b^2 = 8$ with $\sigma_w^2/\sigma_b^2 = (1/3, 1, 3)$, $\mu = (0.5, 1, 2)$, and setting $\rho^2 = \sigma_w^2 + \sigma_b^2 + \mu^2$. We conduct $10^5$ null simulations. Table 1 shows the empirical type I errors for $\sigma_w^2/\sigma_b^2 = 1, \mu = 1$. We compare the proposed test (denoted as QMS) to the Z-score and Z-Wald tests. Overall the type I errors are controlled well for the proposed method, while the Z-score test is very conservative and Z-Wald test has grossly inflated type I errors.

*

Table 1: Empirical type I errors at significance level $\alpha$ estimated over $10^5$ simulations.

| $n = 10$ | | | | | | | | | |
|---|---|---|---|---|---|---|---|---|---|
| | m=5 | | | m=10 | | | m=20 | | |
| $\alpha$ | 0.05 | 0.01 | 0.001 | 0.05 | 0.01 | 0.001 | 0.05 | 0.01 | 0.001 |
| QMS | 0.052 | 0.009 | 2.5e-4 | 0.054 | 0.009 | 5.2e-4 | 0.055 | 0.010 | 7.0e-4 |
| Z-score | 0.004 | 1.0e-4 | 1e-5 | 0.004 | 5e-5 | 0 | 0.004 | 1.4e-4 | 3e-5 |
| Z-Wald | 0.115 | 0.068 | 0.037 | 0.122 | 0.074 | 0.042 | 0.130 | 0.081 | 0.048 |
| $n = 20$ | | | | | | | | | |
| | m=5 | | | m=10 | | | m=20 | | |
| $\alpha$ | 0.05 | 0.01 | 0.001 | 0.05 | 0.01 | 0.001 | 0.05 | 0.01 | 0.001 |
| QMS | 0.052 | 0.009 | 4.6e-4 | 0.053 | 0.010 | 6.1e-4 | 0.053 | 0.010 | 7.1e-4 |
| Z-score | 0.011 | 2.0e-4 | 0 | 0.009 | 1.7e-4 | 0 | 0.010 | 1.7e-4 | 0 |
| Z-Wald | 0.086 | 0.045 | 0.021 | 0.092 | 0.048 | 0.023 | 0.094 | 0.051 | 0.025 |
| $n = 30$ | | | | | | | | | |
| | m=5 | | | m=10 | | | m=20 | | |
| $\alpha$ | 0.05 | 0.01 | 0.001 | 0.05 | 0.01 | 0.001 | 0.05 | 0.01 | 0.001 |
| QMS | 0.051 | 0.009 | 5.4e-4 | 0.052 | 0.010 | 6.3e-4 | 0.051 | 0.010 | 6.4e-4 |
| Z-score | 0.014 | 6.0e-4 | 0 | 0.014 | 4.5e-4 | 0 | 0.013 | 4.3e-4 | 0 |
| Z-Wald | 0.076 | 0.035 | 0.014 | 0.079 | 0.037 | 0.016 | 0.079 | 0.038 | 0.016 |

##### 1.2 Power

We include the proposed test and the Z-score test in the power comparison. We consider the combinations of $n = (10, 20, 30)$, $m = (5, 10, 20)$, $\sigma_w^2 + \sigma_b^2 = 5$ with $\sigma_w^2/\sigma_b^2 = (1/3, 1, 3)$, $\mu = (0.5, 0.75, 1)$, and $\rho = 3$. We are testing $H_0 : \sigma_w^2 + \sigma_b^2 + \mu^2 \geq \rho^2$. Generally larger $(n, m, \sigma_w^2/\sigma_b^2)$, and smaller $\mu$ will lead to larger rejection power. We use $10^4$ Monte Carlo simulations to estimate power under each configuration. Table 2 summarizes the power for $\mu = 0.75$. Overall the proposed method performs better than Z-score test by a large margin. We observed similar patterns for $\mu = 0.5, 1$.

In addition, we also evaluate the power calculation. We consider the same simulation settings as previously and treat the Monte Carlo results as gold standard. We then apply the proposed analytical power calculation methods and compare their results to the gold standard results from the Monte Carlo simulations. Table 3 summarizes the results for $\mu = 0.75$. Overall we have observed that the analytically computed powers (denoted as ACP) are very close to the Monte Carlo results (denoted as MCP).

Table 2: Power (%) at 0.05 significance level.

| $n = 10$ | | | | | | | | | |
|---|---|---|---|---|---|---|---|---|---|
| | $m = 5$ | | | $m = 10$ | | | $m = 20$ | | |
| $\sigma_w^2 : \sigma_b^2$ | 1:3 | 1:1 | 3:1 | 1:3 | 1:1 | 3:1 | 1:3 | 1:1 | 3:1 |
| QMS | 28.8 | 37.2 | 52.5 | 30.3 | 41.3 | 63.3 | 30.5 | 41.9 | 70.0 |
| Z-score | 1.68 | 2.99 | 10.1 | 1.69 | 3.33 | 12.2 | 1.64 | 3.29 | 13.4 |
| $n = 20$ | | | | | | | | | |
| | $m = 5$ | | | $m = 10$ | | | $m = 20$ | | |
| $\sigma_w^2 : \sigma_b^2$ | 1:3 | 1:1 | 3:1 | 1:3 | 1:1 | 3:1 | 1:3 | 1:1 | 3:1 |
| QMS | 49.7 | 65.1 | 82.8 | 51.6 | 69.7 | 91.3 | 53.0 | 71.9 | 95.4 |
| Z-score | 16.6 | 28.1 | 52.5 | 17.6 | 32.2 | 65.4 | 18.2 | 33.8 | 73.9 |
| $n = 30$ | | | | | | | | | |
| | $m = 5$ | | | $m = 10$ | | | $m = 20$ | | |
| $\sigma_w^2 : \sigma_b^2$ | 1:3 | 1:1 | 3:1 | 1:3 | 1:1 | 3:1 | 1:3 | 1:1 | 3:1 |
| QMS | 66.4 | 81.8 | 95.1 | 68.4 | 86.5 | 98.5 | 69.8 | 88.2 | 99.5 |
| Z-score | 35.7 | 54.4 | 81.3 | 37.9 | 61.2 | 91.6 | 38.3 | 63.7 | 95.8 |

### 2 R package

The following are some sample codes to install and use the 'SPprm' R package.

```
## install the package

devtools::install_github('baolinwu/SPprm')

library(SPprm)

## simulate PRM data

n = 20;  m = 10

s2w = s2b = 4;  mu = 1

e = matrix(rnorm(n*m), n,m)*sqrt(s2w)

u = rnorm(n)*sqrt(s2b)

X = mu + u + e

## QMS test

rho = sqrt(s2w+s2b+mu^2)

PRMtest(X, rho)$p.val

PRMtest(X, rho+0.75)$p.val

## power calculation
```

Table 3: Computed power (%) at 0.05 significance level: Monte Carlo results are displayed as MCP, while the analytically computed powers are displayed as ACP.

| $n = 10$ | | | | | | | | | |
|---|---|---|---|---|---|---|---|---|---|
| | $m = 5$ | | | $m = 10$ | | | $m = 20$ | | |
| $\sigma_w^2 : \sigma_b^2$ | 1:3 | 1:1 | 3:1 | 1:3 | 1:1 | 3:1 | 1:3 | 1:1 | 3:1 |
| MCP | 28.8 | 37.2 | 52.5 | 30.3 | 41.3 | 63.3 | 30.5 | 41.9 | 70.0 |
| ACP | 26.8 | 34.9 | 48.5 | 27.7 | 38.2 | 58.1 | 28.2 | 40.1 | 64.7 |
| $n = 20$ | | | | | | | | | |
| | $m = 5$ | | | $m = 10$ | | | $m = 20$ | | |
| $\sigma_w^2 : \sigma_b^2$ | 1:3 | 1:1 | 3:1 | 1:3 | 1:1 | 3:1 | 1:3 | 1:1 | 3:1 |
| MCP | 49.7 | 65.1 | 82.8 | 51.6 | 69.7 | 91.3 | 53.0 | 71.9 | 95.4 |
| ACP | 48.6 | 62.6 | 80.2 | 50.3 | 67.7 | 89.3 | 51.1 | 70.5 | 93.7 |
| $n = 30$ | | | | | | | | | |
| | $m = 5$ | | | $m = 10$ | | | $m = 20$ | | |
| $\sigma_w^2 : \sigma_b^2$ | 1:3 | 1:1 | 3:1 | 1:3 | 1:1 | 3:1 | 1:3 | 1:1 | 3:1 |
| MCP | 66.4 | 81.8 | 95.1 | 68.4 | 86.5 | 98.5 | 69.8 | 88.2 | 99.5 |
| ACP | 66.0 | 80.7 | 93.6 | 67.9 | 85.1 | 97.9 | 68.8 | 87.3 | 99.2 |

```
## analytical calc

PRMap(alpha=0.05,n, m, s2w, s2b, mu, rho+0.75)$pwr

## Monte Carlo sim

PRMmcp(alpha=0.05,n, m, s2w, s2b, mu, rho+0.75)$pwr
```
